# Detection of succinate by intestinal tuft cells triggers a type 2 innate immune circuit

**DOI:** 10.1101/310110

**Authors:** Marija S. Nadjsombati, John W. McGinty, Miranda R. Lyons-Cohen, Joshua L. Pollack, G.A. Nagana Gowda, David J. Erle, Richard M. Locksley, Daniel Raftery, Jakob von Moltke

## Abstract

Initiation of immune responses requires innate immune sensing, but immune detection of the helminths, protists, and allergens that stimulate type 2 immunity remains poorly understood. In the small intestine, type 2 immune responses are regulated by a tuft cell-ILC2 signaling circuit. Tuft cells express components of a canonical taste transduction pathway, including the membrane channel TRPM5, but the ligands and receptors that activate tuft cells in the small intestine are unknown. Here we identify succinate as the first ligand that activates intestinal tuft cells to initiate type 2 immune responses. Using mRNA-Seq on tuft cells from different tissues, we show that all tuft cells express the intracellular taste transduction pathway, but expression of upstream receptors is tissue-specific. In the small intestine, tuft cells express the succinate receptor SUCNR1. Remarkably, providing succinate in drinking water is sufficient to induce a multifaceted type 2 immune response in the murine small intestine, involving all known components of the tuft-ILC2 circuit. The helminth *Nippostrongylus brasiliensis* secretes succinate as a metabolite, and sensing of both succinate and *N. brasiliensis* requires tuft cells and TRPM5, suggesting a novel paradigm in which type 2 immunity monitors microbial metabolism. Manipulation of succinate sensing may have therapeutic benefit in numerous intestinal diseases.

## Introduction

Innate immune sensing, involving the binding of microbially derived ligands to host receptors, is the fundamental first step in initiation of immune responses to microbes. The ligand-receptor paradigm of innate immune sensing was first proposed by Charles Janeway (Janeway, 1989), and soon borne out experimentally by the discovery of toll-like receptor 4, which binds bacterial lipopolysaccharide. This seminal advance ushered in an era of rapid progress in the newly established field of innate immune sensing. In the last two decades, many more innate immune receptors have been discovered, virtually all of which recognize conserved ligands derived from viruses, bacteria, or fungi. As a result, our understanding of innate immune detection for these classes of pathogens is quite advanced. However, much less is known about innate immune sensing of helminths, intestinal protists, and allergens, all of which can induce a type 2 immune response defined by the production of interleukin(IL)-4, −5, −9, and −13. These canonical type 2 cytokines activate and recruit type 2 leukocytes, such as basophils, eosinophils, mast cells, alternatively activated macrophages, and type 2 helper T cells (Th2). IL-13 in particular also targets non-hematopoietic tissues, leading to profound changes in tissue physiology, such as epithelial remodeling and hypercontractility of smooth muscle. In the case of helminth infection, these coordinated responses are designed to expel or at least restrict and wall off the worms. When activated chronically by exposure to allergens, type 2 responses result in asthma and fibrosis.

In investigating the initiation of type 2 immune responses, there is currently much interest in the activation and function of group 2 innate lymphoid cells (ILC2s). ILC2s are the dominant innate source of IL-5, −13, and −9 in numerous models of type 2 inflammation (Halim, 2016), and unlike most other type 2 immune cells, ILC2s are tissue-resident, which could poise them as innate immune sentinels. While ILC2s are indeed among the first type 2 cells activated, there is no evidence that ILC2s sense helminths or other type 2 stimuli directly; instead, they are activated by host-derived signals from the surrounding tissue (von Moltke and Locksley, 2014). ILC2 activating signals include the cytokines IL-25, TSLP, IL-33, and TL1A, as well as leukotrienes and neuronal peptides (Cardoso et al., 2017; Klose and Artis, 2016; Klose et al., 2017; Wallrapp et al., 2017). The relative contribution of these signals to ILC2 activation varies by tissue, and ILC2s must integrate multiple signals for optimal activation (von Moltke et al., 2017).

In the small intestine, IL-25 is critical for ILC2 activation and worm clearance (Fallon et al., 2006; Neill et al., 2010). Recently, we and others demonstrated that IL-25 is made exclusively by epithelial tuft cells, which drive a feed-forward immune signaling circuit that is required for innate and perhaps adaptive type 2 responses in the small intestine (Gerbe et al., 2016; Howitt et al., 2016; von Moltke et al., 2016). In this circuit, tuft cell-derived IL-25 induces IL-13 production by ILC2s. IL-13 then signals in undifferentiated epithelial progenitors, biasing their lineage commitment towards goblet and tuft cells, the latter of which further promote ILC2 activation, resulting in the feed-forward loop that we will refer to as the tuft-ILC2 circuit. Since the entire intestinal epithelium is replaced every 5–7 days (Barker, 2014), the induction of IL-13 rapidly leads to epithelial remodeling marked by goblet and tuft cell hyperplasia. Furthermore, production of IL-13, −5, and −9 by intestinal ILC2s promotes eosinophilia, alternative activation of macrophages, and other hallmarks of a type 2 immune response.

In most specific pathogen-free mice, activation of the tuft-ILC2 circuit is restrained and tuft cells are extremely rare, representing <1% of all epithelial cells (Gerbe et al., 2016). However, the protist *Tritrichamonas muris*, a parabasalid that exists in the natural flora of some vivariums (Chudnovskiy et al., 2016; Escalante et al., 2016), is specifically known to activate the tuft-ILC2 circuit and downstream type 2 responses (Howitt et al., 2016). In addition, infection with helminths such as the rat-adapted hookworm *Nippostrongylus brasiliensis* leads to even greater activation of the tuft-ILC2 circuit, and in this context tuft cell frequency increases about 10 fold within 7–9 days, corresponding to the time frame within which worms are cleared (von Moltke et al., 2016). In sum, tuft cells are a recently identified component of type 2 immunity and drive a signaling circuit that initiates the canonical intestinal type 2 immune response. How the circuit is activated, however, remains completely unknown.

Tuft cells were first discovered in the 1950s, when the advent of electron microscopy allowed researchers to observe their unique morphology, defined by the presence of an apical bundle of long microvilli (Isomäki, 1962; Jarvi and Keyrilainen, 1956). Tuft cells have been identified in the epithelia of numerous tissues of mice and rats, including the entire intestinal tract, the gall bladder, pancreatic ducts, the upper airways, the urethra, and the thymus (Akimori et al., 2011; Deckmann et al., 2014; Finger et al., 2003; Krasteva et al., 2011; Luciano and Reale, 1997; Panneck et al., 2014). Cells with tuft-like morphology and marker expression have also been noted in human airways and intestine, but remain poorly characterized, especially in the intestine (Braun et al., 2011; DiMaio et al., 1988; Lee et al., 2014; Moxey and TRIER, 1978).

In 2008, Damak and colleagues provided the first transcriptional analysis of murine intestinal tuft cells, and confirmed earlier suggestions that tuft cells encode a chemosensing pathway that was first described in the context of taste transduction (Bezençon et al.,

2008). In taste buds, surface-expressed G protein-coupled taste receptors bind their ligands and activate a specialized alpha subunit called a-gustducin, encoded by the gene *Gnat3* (Chaudhari and Roper, 2010). Gustducin in turn activates phospholipase C beta 2 (PLCB2) leading to the production of IP3, release of intracellular Ca^2+^ stores, and sodium influx via the calcium-gated transient receptor potential cation channel subfamily M member 5 (TRPM5). The resulting depolarization of the cell drives release of neuromediators that activate nearby nerve terminals. Various components of this chemosensing pathway have been reported in tuft cells of numerous tissues, and, importantly, a *Trpm5-GFP* reporter mouse demonstrated that expression of *Trpm5* in the intestinal tract is restricted to tuft cells (Bezençon et al., 2008). The transcriptional analysis of intestinal tuft cells also revealed that they express several genes previously associated only with neurons, such as the microtubule-associated doublecortin-like kinase 1 (DCLK1), which was later shown to be a definitive marker of intestinal tuft cells (Gerbe et al., 2009). Further, tuft cells and some taste cell lineages share a requirement for the transcription factor POU domain, class 2, transcription factor 3 (POU2F3) and these cells are all absent in *Pou2f3^-/-^* mice (Gerbe et al., 2016; Matsumoto et al., 2011).

Since expression of the TRPM5-dependent chemosensing pathway was first noted, evidence of a sensing function for tuft cells has mounted. Tuft cells of the airways and urethra express a subset of canonical taste receptors and have been shown to regulate smooth muscle contractions in response to bitter ligands and bacterial quorum-sensing molecules (Deckmann et al., 2014; Krasteva et al., 2011; Tizzano et al., 2010). In the intestine, the tuft cell hyperplasia induced by *T. muris* was shown to require *Trpm5* and *Gnat3* (Howitt et al., 2016). Therefore, it has been widely postulated that tuft cells are immune sentinels, but a ligand(s) and receptor(s) that activate intestinal tuft cells and the tuft-ILC2 circuit remain unknown.

Here we show that the metabolite succinate is sensed by tuft cells in the small intestine and is sufficient to induce a type 2 immune response *in vivo*. First, using mRNA sequencing of tuft cells from five different tissues, we identify a core tuft cell signature that includes intracellular components of the TRPM5 chemosensing pathway, but strikingly lacks initiating chemoreceptors, suggesting that tuft cells employ different surface receptors to match the sensing requirements of each tissue. In the small intestine, tuft cells selectively express the G protein coupled receptors *Ffar3*, a short chain fatty acid (SCFA) receptor, and *Sucnr1*, the receptor for extracellular succinate. Remarkably, dietary supplementation with succinate, but not SCFAs, is sufficient to induce a robust and multifaceted type 2 immune response in the murine small intestine. This response requires TRPM5 and all components of the tuft-ILC2 circuit, including IL-25, ILC2s, and IL-13. Additionally, we find *N. brasiliensis* secretes succinate as a metabolic waste product and can activate SUCNR1 in heterologous cells. The immune sensing of both succinate and *N. brasiliensis* requires both TRPM5 and tuft cells, suggesting that sensing of succinate contributes to the type 2 immune response elicited by helminth infection. In sum, our findings identify the first intestinal tuft cell ligand and its cognate receptor. More broadly, our results define metabolite sensing as a novel paradigm in the initiation of type 2 immunity.

## Results

### mRNA sequencing identifies a transcriptional tuft cell signature

To identify both the unique and shared features of tuft cells in different tissues, we used IL-25 reporter mice (Flare25; von Moltke et al., 2016) to sort-purify CD45^lo^ EPCAM^+^ RFP^+^ tuft cells from the small intestine, gall bladder, colon, thymus, and trachea for mRNA sequencing. We also sorted CD45^lo^ EPCAM^+^ RFP^-^ cells from the small intestine to serve as a non-tuft epithelial reference. Principal component analysis (PCA) of normalized mRNA sequencing read counts clustered tuft cells from each tissue separately and also identified related features in tuft cells from the intestinal tract (gall bladder, small intestine, and colon) compared to those from the thymus and trachea (Figure 1A). The tissue-specific signature of each tuft cell population was also identifiable following hierarchical clustering of all differentially expressed genes (FDR < 0.01; absolute fold change > 8) (Figure 1B, Figure S1, and Item S1). Interestingly, tuft cells from the thymus had the greatest number of differentially expressed genes and clustered farthest from other tuft cells by PCA. The thymus is the only immune organ known to contain tuft cells and also the only tissue in which tuft cells are not found in an epithelial barrier surrounding a hollow lumen. It is likely therefore that thymic tuft cells have unique functions, perhaps related to T cell development and education.

**Figure 1.**
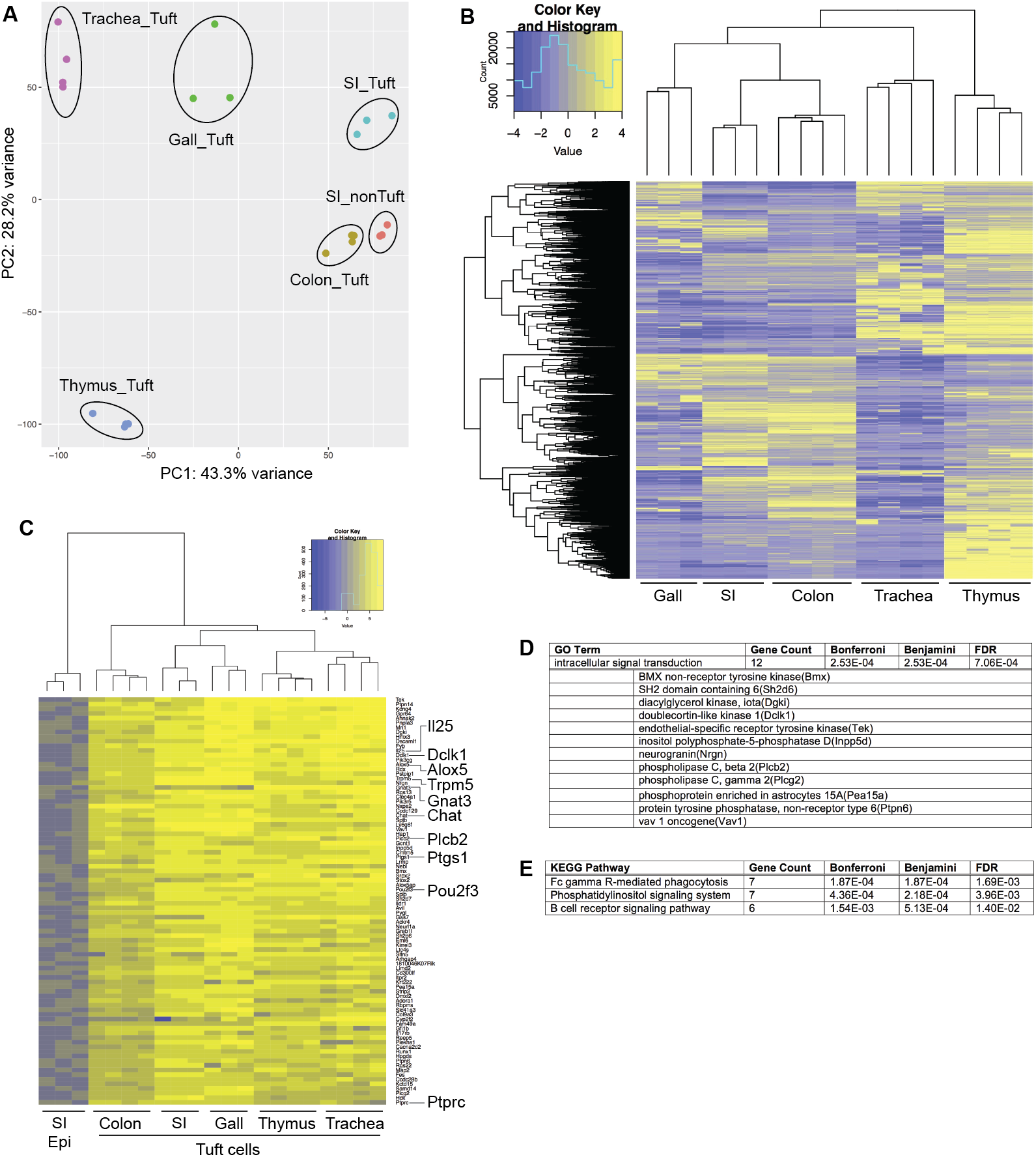
RNA-Seq identifies a transcriptional tuft cell signature. Tuft cells (CD45^lo^ EPCAM^+^ Flare25^+^) were sorted from the indicated tissues of naïve B6._*I125/Flare25/Flare2S*_ mice for mRNA sequencing. Non-tuft epithelial cells of the small intestine (SI; CD45^lo^ EPCAM^+^ Flare25-) were sorted as negative controls. **(A)** Principle component analysis of gene expression data. Each dot represents an individual mouse, except in samples from the trachea where each dot represents cells pooled from four mice. **(B)** Hierarchical clustering of differentially expressed genes among tuft cell subsets with an absolute fold change > 8 (FDR < 0.01]. **(C)** Ranked listing of a tuft cell transcriptional signature, comprised of genes that were expressed Log2 fold-change > 4 in all tuft cells relative to non-tuft epithelial cells. **(D)** GO Term enrichment analysis identified a single overrepresented gene category within the tuft cell signature (FDR < 0.05). **(E)** Overrepresented pathways identified by KEGG analysis (FDR < 0.05).

Next we used the mRNA sequencing data to define a transcriptional signature that is common to tuft cells across all tissues analyzed (Figure 1C and Supplementary Item 2). As expected, this signature included *Il25* and *Pou2f3*, the transcriptional regulator required for specification of the tuft cell lineage (Gerbe et al., 2016). Other tuft cell markers that were previously characterized in individual tissues, such as *Dclkl, Ptgsl, Alox5, Ptprc* (CD45), and *Chat*, were also included in our global tuft cell signature. To analyze this signature further, we performed GO Term and KEGG pathway enrichment analysis. Interestingly, the only significantly enriched GO Term among tuft cell signature genes was “intracellular signal transduction” (FDR < 0.05; Figure 1D and Item S3), which includes genes like *Plcb2, Plcg2, Nrgn*, and *Dclk1*. KEGG pathway analysis further underscored the signaling capacity of tuft cells (Figure 1E and Item S4).

### The receptor repertoire of tuft cells is tissue-specific

Next, we specifically interrogated expression of the TRPM5-dependent chemosensing pathway that has been implicated in tuft cell function and found the same pattern observed in the global tuft cell signature. Downstream intracellular components of the chemosensing pathway, such as *Plcb2* and *Trpm5*, are included in the tuft cell signature and are expressed across all tuft cell subsets (Figures 1C and 2A, B). *Gnat3*, the G alpha subunit that couples to taste receptors, also appears in the global signature but its expression is more variable across tissues (Figures 1C and 2C). Canonical taste receptors, meanwhile, are completely absent in the tuft cell signature (Figure 2D). This lack of receptors was at first unexpected given the putative sensory function of tuft cells, but suggests an intriguing model. We hypothesized that the core intracellular machinery of signal transduction is conserved across all tuft cells but that the receptors that activate these pathways are differentially expressed depending on the sensing requirements of each tissue. Indeed, as has been previously reported, a subset of canonical taste receptors (TasR) is expressed in tracheal tuft cells (Krasteva et al., 2011 and Figure 2D). We also detected expression of taste receptors in thymic tuft cells, but they are absent in intestinal tuft cells, especially those of the small intestine.

**Figure 2.**
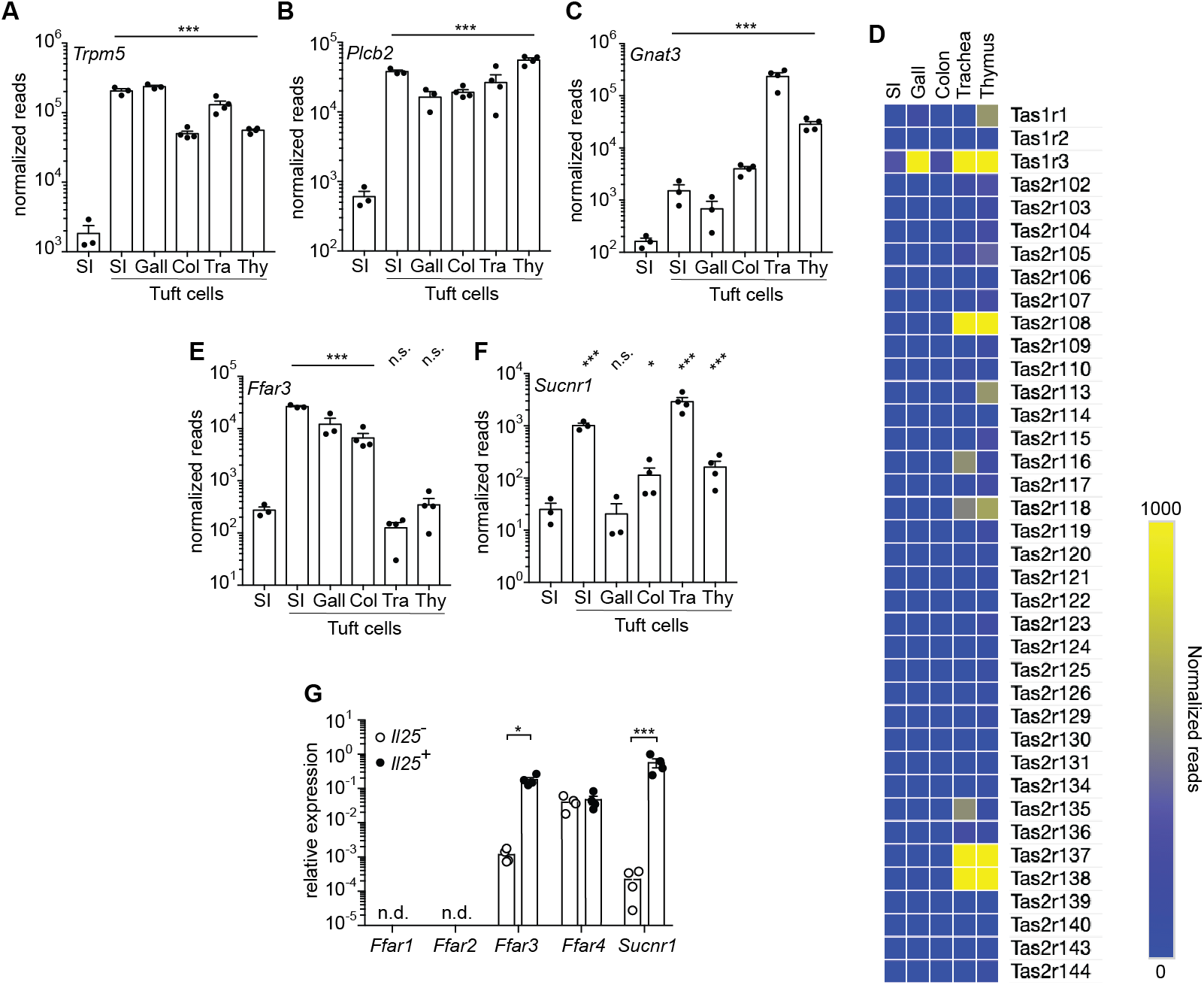
The chemosensory receptor repertoire of tuft cells is tissue-specific (A-C, E, F) RNA-Seq data generated as described in Figure 1 were analyzed for expression of the indicated genes in tuft cells sorted from small intestine (SI), gall bladder (Gall), colon (Col), trachea (Tra), and thymus (Thy). Non-tuft small intestinal epithelium served as a negative control. (D) Heat map depicting expression of all taste receptors in tuft cells from the indicated tissues. (G) Tuft cells (I125^+^) and non-tuft cells (1125’) were sorted from the small intestine of naïve B6.*I125^Flare25/FIare25^* mice and expression of the indicated genes was assessed by RT-qPCR. A-F show biological replicates from one RNA sequencing experiment. Data in G show biological replicates pooled from three experiments. In A-C and E-G, mean + SEM is shown. In A-C and E-F *, FDR < .05; ***, FDR < .001 by statistical analysis in RNA sequencing pipeline. In G *, p < 0.05; ***, p < 0.001 by multiple t-tests (G). n.s., not significant.

**Figure S1.**
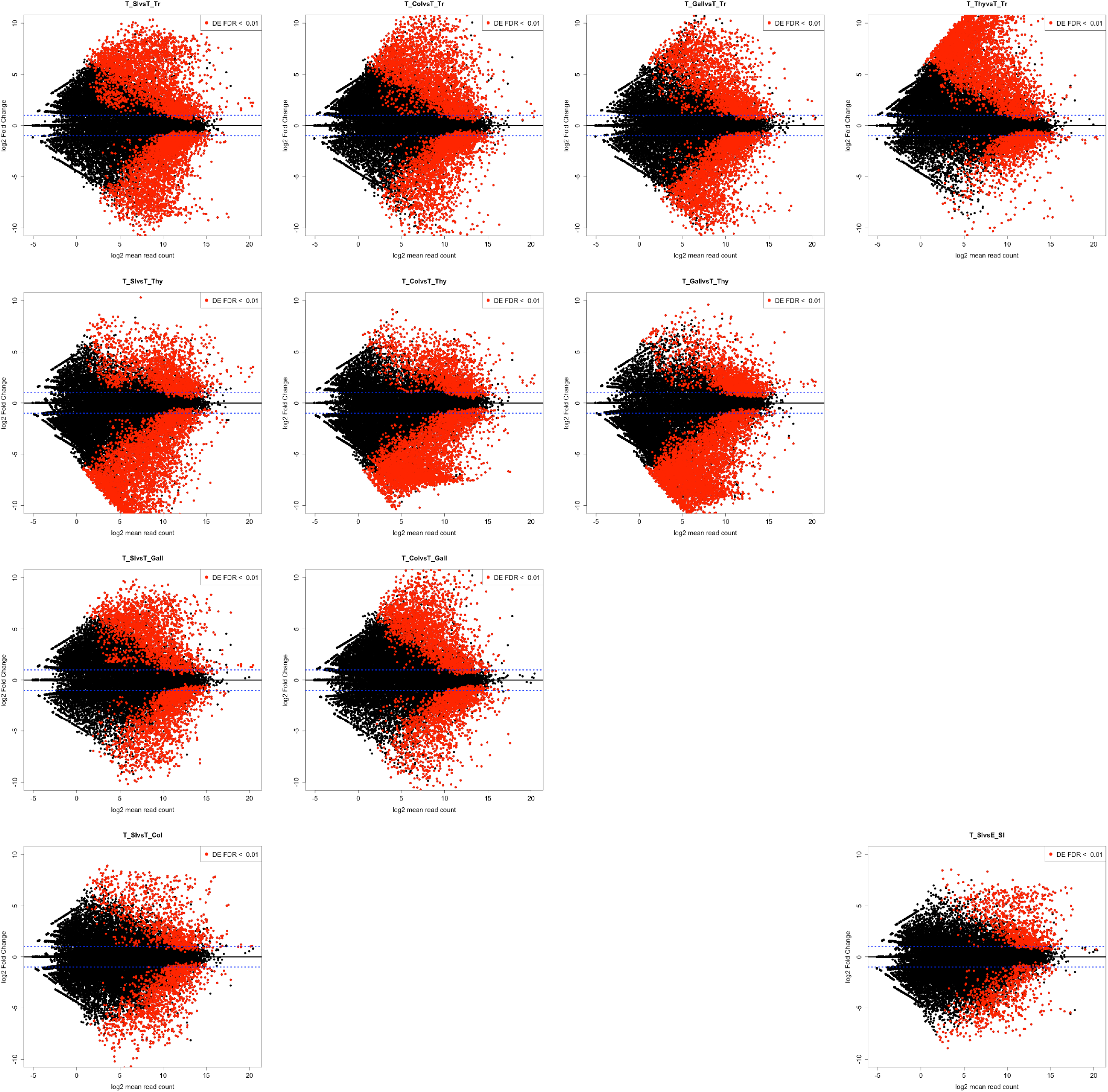
**I** Volcano plots showing pairwise comparisons of differentially expressed genes in the cell subsets analyzed by mRNA sequencing. Red dots represent genes with FDR < 0.01. T_ = tuft cells; E_ = non-tuft epithelium; SI = small intestine; Col = colon; Gall = gall bladder; Thy = thymus; Tr = trachea

Since the immune function of tuft cells is best characterized in the small intestine, we sought to identify the receptors required for chemosensing in these cells. Taste receptors are GPCRs and TRPM5 activation is Ca^2+^-dependent; therefore we began by searching for cell surface GPCRs that might induce Ca^2+^ flux and that are highly and selectively expressed in small intestinal tuft cells. Interestingly, two metabolite sensors match these criteria. FFAR3 is the receptor for the short chain fatty acids (SCFA) propionate and butyrate, and has been implicated in sensing these bacterial metabolites in the intestine (Trompette et al., 2014). SUCNR1 is the extracellular receptor for the citric acid cycle intermediate succinate and is best characterized in the kidney, where it helps to regulate blood pressure (Ariza et al., 2012). mRNA sequencing results showed that *Ffar3* is enriched in tuft cells of the intestinal tract, while *Sucnr1* is expressed primarily in tuft cells of the small intestine and trachea (Figure 2E, F). To validate these sequencing results, we again sorted tuft cells (CD45^lo^ Epcam^+^ Flare25^+^) and non-tuft epithelial cells (CD45^lo^ Epcam^+^ Flare25^-^) from the small intestine and performed qPCR to confirm that both *Ffar3* and *Sucnr1* are highly expressed and enriched in small intestinal tuft cells (Figure 2G). Other members of the SCFA receptor family were either undetectable *(Ffar1, Ffar2)* or expressed equally in tuft cells and non-tuft epithelium *(Ffar4)*. SUCNR1 and FFAR3 therefore warranted further analysis as candidate immune receptors in tuft cells.

### Succinate is sufficient to induce a type 2 immune response in the small intestine

To test whether tuft cells can sense lumenal SCFA and/or succinate, we supplemented mouse drinking water with 150 mM (Na^+^)_2_-succinate, Na^+^-propionate, Na^+^-acetate, Na^+^-butyrate, or NaCl and measured tuft cell hyperplasia as a readout for the intestinal type 2 immune response. Remarkably, succinate, but not SCFAs or NaCl, was sufficient to induce a >5-fold increase in tuft cells in the small intestine (Figure 3A, B). Although tuft cell hyperplasia was detectable throughout the small intestine, it was most pronounced in the distal segment (terminal 10 cm), which we used for further characterization (Figure 3C). The succinate-induced tuft cell hyperplasia was maximal at a concentration of 150 mM given ad libitum in drinking water. Concentrations of 75 mM and 300 mM both elicited less tuft cell hyperplasia, the latter likely due to reduced water consumption at such high salt concentrations (Figure 3D). Kinetically, an increase in tuft cell numbers was detectable just two days after administering succinate, but continued to rise until day 7-8 (Figure 3E). We noted a downward trend in tuft cell numbers at later timepoints, perhaps due to regulatory feedback mechanisms, but tuft cell frequency remained well above background throughout the time course. Unless otherwise noted, further experiments were performed by analyzing the distal (last 10 cm) small intestine after 7 days of 150 mM succinate administration. Of note, dietary succinate was sufficient to induce multiple additional components of the canonical intestinal type 2 immune response, including goblet cell hyperplasia and hypertrophy (Figure 3F-H). In the lamina propria, ILC2 activation (measured using the Smart13 reporter of IL-13 production) was evident just 36 hours after initiating succinate treatment (Figure 3I-J and Figure S3A), and after 7 days we found further ILC2 activation and accumulation in the mesenteric lymph nodes (MLN) (Figure 3K-M and Figure S3B). ILC2 activation in the MLN was accompanied by recruitment of eosinophils, another hallmark of the type 2 immune response (Figure 3N).

**Figure 3.**
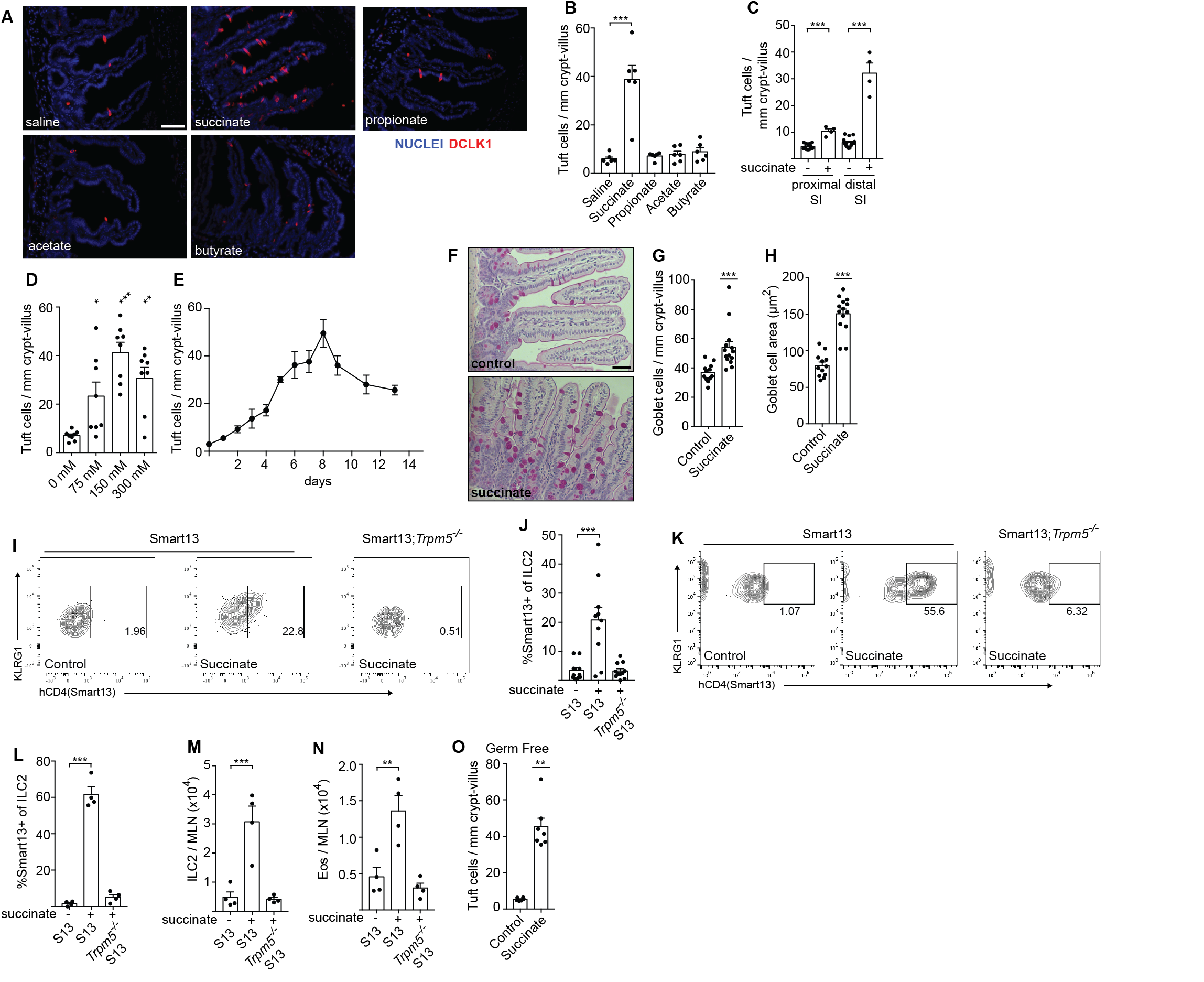
Succinate is sufficient to induce a type 2 immune response in the small intestine. **(A-C)** Mice were given the indicated molecules (150 mM, except 300 mM for NaCl) in drinking water for 7 d prior to harvest, sectioning, and labeling of small intestine with anti-DCLK1 (tuft cell label; red) and DAPI (blue). **(A)** Representative images of distal (last 10 cm) small intestine are shown. Scale bar = 50 μm **(B)** Tuft cells were quantified by microscopy. **(C)** Tuft cells were enumerated in the proximal (first 10 cm) and distal (last 10 cm) sections of the small intestine (SI) of mice treated 7 d with 150 mM succinate or a water control. **(D)** Tuft cells were enumerated in the distal (last 10 cm) small intestine following 7 d treatment with the indicated concentrations of succinate. **(E)** Kinetics of tuft cell expansion in distal (last 10 cm) small intestine of mice treated with 150mM succinate (n = 3-9 biological replicates for each timepoint). **(F)** Representative images of middle (10-20 cm from cecum) small intestine stained with periodic acid-Schiff to visualize goblet cell number and hypertrophy following 7 d of 150 mM succinate treatment. Scale bar = 50 μm. Goblet cell numbers **(G)** and hypertrophy **(H)** were quantified by microscopy. **(I)** ILC2 activation was assessed in the lamina propria after 36 hours of 150mM succinate treatment of wild-type or *Trpm5-/-* IL-13 reporter mice (Smartl3). Representative flow plots are shown. **(J)** Quantification of ILC2 activation. **(K-N)** Mesenteric lymph node cells were collected after 7 d treatment with 150 mM succinate and assessed for ILC2 production of IL-13 **(K, L)** and absolute numbers of ILC2s **(M)** and eosinophils **(N)**. **(O)** Tuft cell expansion was measured in distal (last 10 cm) small intestine of germ-free mice after 7 d treatment with 150 mM succinate. All bar graphs depict mean + SEM of at least two independent experiments except that K-N depict results from one experiment. Each symbol represents an individual mouse. *, p < 0.05; **, p < 0.01; ***, p < 0.001 by one-way ANOVA (B-D, J, L-N) with comparison to control, or by Mann-Whitney (G-H, O).

Since commensal bacteria both produce and consume succinate(Ferreyra et al., 2014; De Vadder et al., 2016), it is possible that dietary succinate supplementation alters the composition of the intestinal microbiome and activates tuft cells indirectly. To confirm that succinate can induce tuft cell hyperplasia in the absence of commensal microbes, we provided succinate in the drinking water of germ free mice and found that succinate-induced tuft cell hyperplasia was equivalent in germ free and SPF mice (Figure 3O). Taken together, our data demonstrate that succinate is an innate immune ligand and is sufficient to induce a multifactorial innate type 2 immune response in the small intestine.

**Figure S2.**
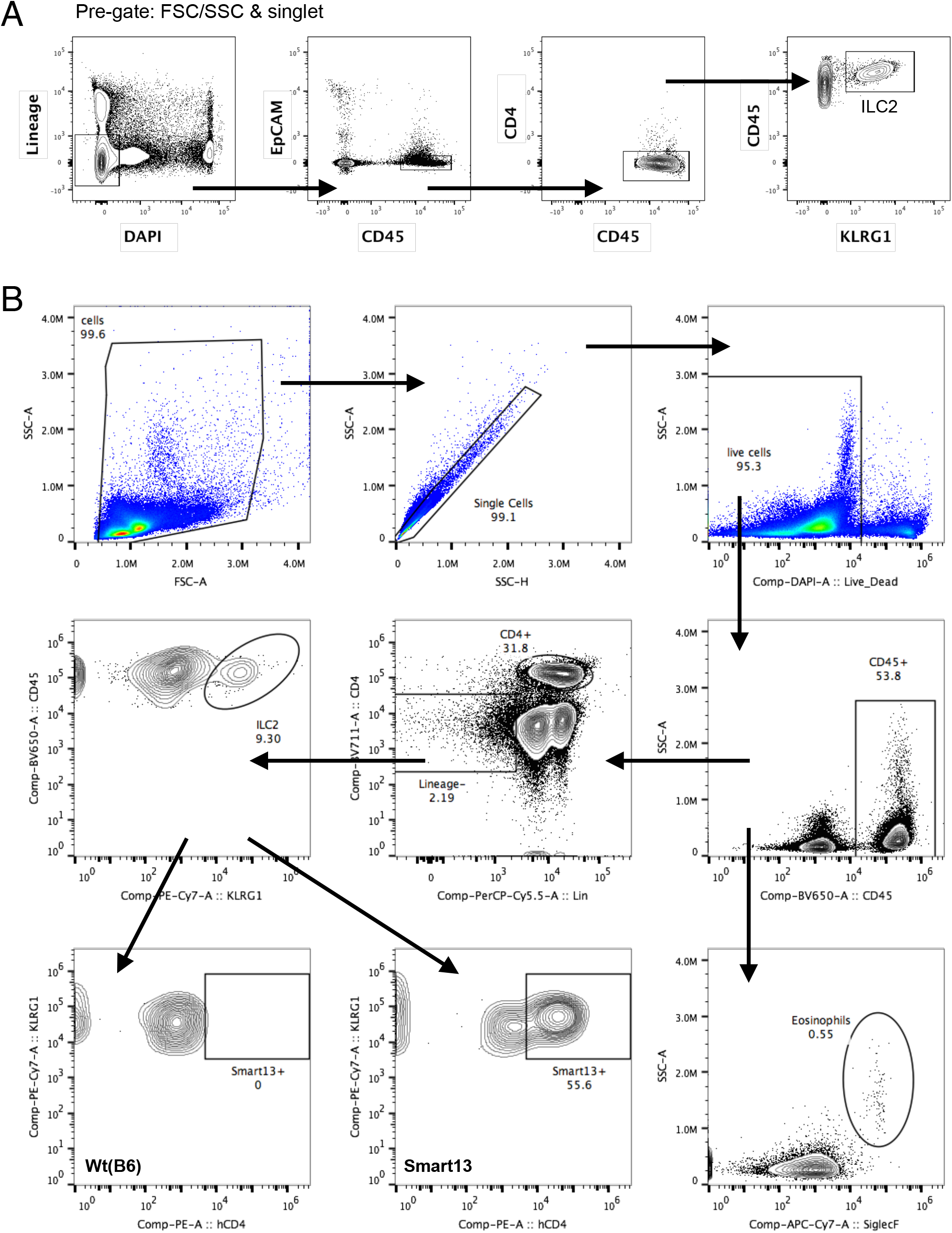
**I** Flow cytometry gating strategy for cells from the small intestine lamina propria (A) and mesenteric lymph nodes (B)

**Figure S3.**
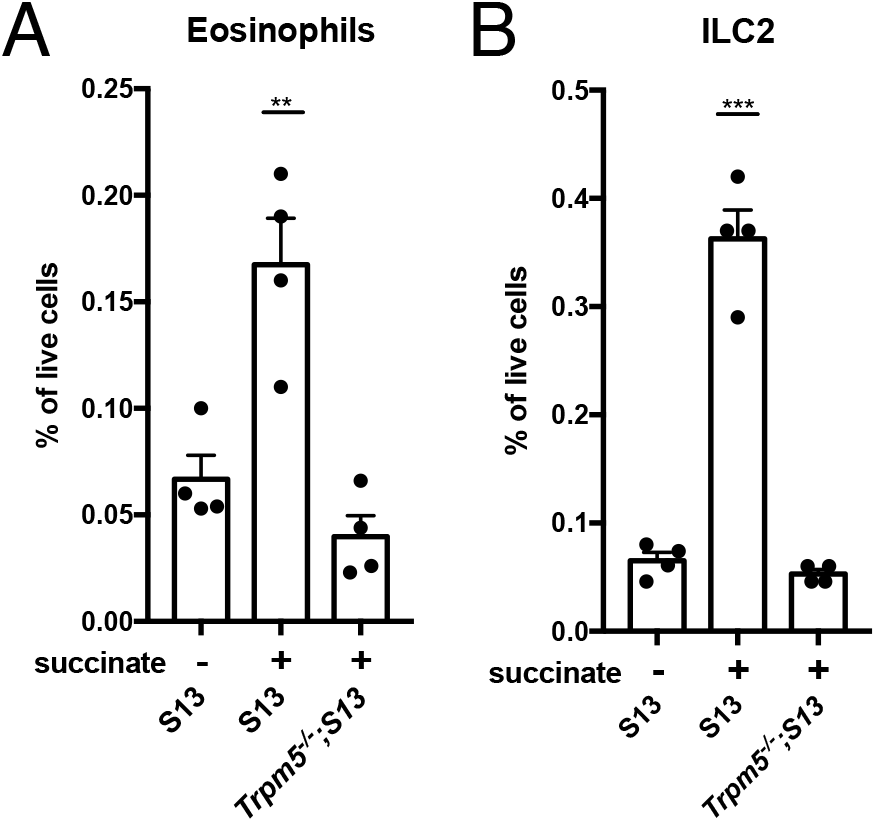
**I** Smart13 or *Trpm5-/-*;Smart13 mice were either left untreated or given 150 mM succinate in the drinking water for 7 days before mesenteric lymph nodes were harvested and analyzed by flow cytometry. (A) Percentage of eosinophils among all live cells. (B) Percentage of ILC2s among all live cells. Graphs depict mean + SEM. Each symbol represents an individual mouse and data are from one experiment. **, p < .01; ***, p < . 001 by one-way ANOVA with comparison to untreated Smart13 mice.

### Succinate signals via the tuft cell-ILC2 circuit

To further characterize the signaling cascade through which dietary succinate is sensed and induces tuft cell hyperplasia, we employed mice deficient in various components of the tuft-ILC2 circuit that regulates innate type 2 immunity in the small intestine. As in helminth infection, succinate-induced tuft cell hyperplasia was completely dependent on IL-4 receptor alpha *(Il4ra)*, likely due to the established requirement for IL-13 signaling in epithelial stem cells to induce tuft and goblet cell hyperplasia (Figure 4A, B). ILC2s are the dominant innate source of IL-13; accordingly, *Il2rg^-/-^* mice, which are broadly lymphopenic, completely failed to induce tuft cell hyperplasia in response to succinate, while the response was largely intact, although still significantly different, in *Rag1^-/-^* mice lacking only adaptive lymphocytes. Tuft cells are the exclusive source of the ILC2-activating cytokine IL-25 in the small intestine, and tuft cell hyperplasia was completely absent in *Il25^-/-^* mice treated with succinate. IL-33, another ILC2-activating cytokine that signals through ST2 but is not expressed in tuft cells (von Moltke et al., 2016), is not required for this response. Lastly, succinate-induced tuft cell hyperplasia, rapid ILC2 activation, and responses in the MLN all require TRPM5 (Figures 4A, B and 3I-N). Since both *Sucnr1* and *Trpm5* are expressed exclusively in tuft cells in the small intestinal epithelium, this last finding strongly suggests that succinate sensing occurs directly in tuft cells, likely via SUCNR1-induced Ca^2+^ flux. Accordingly, goblet cell hyperplasia and hypertrophy are absent in tuft cell-deficient *Pou2f3^-/-^* mice given succinate (Figure 4C-E). Taken together, our data demonstrate that succinate-induced activation of TRPM5 in tuft cells activates the feed-forward tuft-ILC2 circuit to drive an intestinal type 2 immune response.

**Figure 4.**
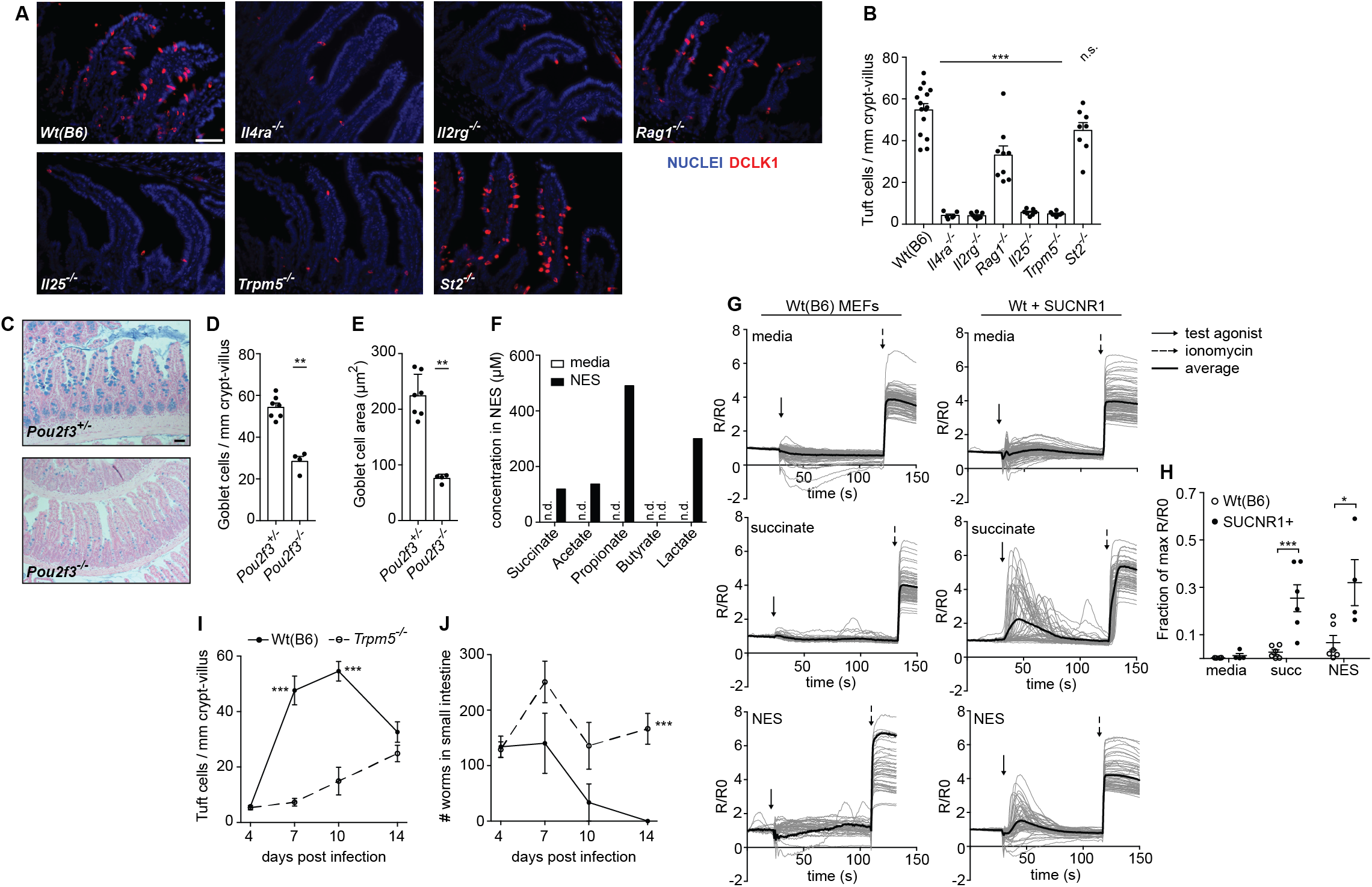
Succinate signals via the tuft cell-ILC2 circuit in a TRPM5-dependent manner. **(A-C)** Distal (last 10 cm) small intestine was harvested from mice of the indicated genotypes after 7 d treatment with 150 mM succinate and labeled with anti-DCLKl (tuft cell label; red) and DAPI (blue). **(A)** Representative images are shown. Scale bar = 50 μm **(B)** Tuft cells were quantified by microscopy. **(C)** Representative Alcian blue staining in middle (10-20 cm from cecum) small intestine from tuft cell-replete (*Pou2/3*^+/-^) or tuft cell-deficient (*Pou2f3*^-/-^) mice following 7 d treatment with 150 mM succinate. Scale bar = 100 μm. Goblet cell numbers **(D)** and hypertrophy **(E)** were quantified by microscopy. **(F)** Concentration of the indicated molecules was measured in *N. brasiliensis* excretory-secretory product (NES) or culture media control by NMR. Results shown are representative of three independent experiments. **(G)** Wild-type (B6) or SUCNR1-transduced MEFs were loaded with a fluorescent calcium indicator and fluorescence/background (R/R_0_) was quantified overtime. Each gray trace represents one cell. Black line indicates average R/R_0_. Arrows indicate time of addition of test agonist or ionomycin positive control (lug/mL). **(H)** The maximal R/Ro of the test agonist was normalized to the R/Ro maximal response to ionomycin for each fluorescent calcium flux sample and quantified. **(I)** Wild-type and *Trpm5-/-* mice were infected with *N. brasiliensis* and tuft cell frequency was assessed in the distal (last 10 cm) small intestine over time. **(J)** Total worm burden in wild-type or TRPM5-deficient mice over the course of *N. brasiliensis* infection. In B, D, and E, each symbol represents one mouse pooled from at least two independent experiments. I and J show the average of 3-10 biological replicates pooled from three independent experiments. G shows representative graphs and H shows technical replicates from more than three independent experiments. All graphs depict mean + SEM. *, p < 0.05; **, p < 0.01; ***, p < 0.001 by one-way ANOVA (B) with comparison to Wt(B6) control, by Mann-Whitney (D-E), or using multiple t-tests (H-J). n.s., not significant; n.d., not detectable.

### *N. brasiliensis* secretes succinate and induces a TRPM5-dependent immune response

Given the link between tuft cells and the type 2 immune response, we hypothesize that succinate sensing is a mechanism for monitoring microbial metabolism in the intestinal lumen. When the cells of land vertebrates are deprived of oxygen, they switch to fermentative metabolism, generating and secreting lactate by using pyruvate as a terminal electron acceptor. In contrast, in helminths, protists, and bacteria, the fermentative metabolic pathways are far more diverse and can lead to the secretion of hydrogen, ethanol, acetate, propionate, lactate, and other metabolites (Müller et al., 2012). Importantly, numerous bacterial species, some protists (e.g. *Tritrichomonas foetus)*, and some helminths (Müller et al., 2012; Tielens, 1994) are known to use fumarate as a terminal electron acceptor during fermentation, resulting in production and release of succinate.

Given the ability of succinate to induce an intestinal immune response with all the hallmarks of a helminth infection, we wondered if succinate sensing by tuft cells might contribute to the immune response to *N. brasiliensis*. A study from 1971 reported that *N. brasiliensis* do indeed secrete succinate, but this has not been further investigated (Saz et al., 1971). To more fully characterize *N. brasiliensis* metabolites, we analyzed *N. brasiliensis* excretory-secretory product (NES) using nuclear magnetic resonance (NMR), and found that acetate, propionate, lactate, and succinate, but not butyrate, are all produced (Figure 4F). Furthermore, NES induced Ca^2+^ flux in mouse embryonic fibroblasts transduced with *Sucnr1*, but not in untransduced cells (Figure 4G, H). We attempted to measure succinate levels in the small intestine of *N. brasiliensis-infected* mice, but even in mice given 150 mM succinate in the drinking water we could not reliably detect increased succinate levels in the proximal or distal small intestine (data not shown), likely because succinate absorption is very rapid in this organ (De Vadder et al., 2016; Wolffram et al., 1994). Thus, our data show that *N. brasiliensis* produces succinate that can induce the Ca^2+^ flux required to open TRPM5 and stimulate downstream signaling.

Lastly, we tested if sensing of *N. brasiliensis* requires TRPM5, as it does for *T. muris* and succinate. Strikingly, tuft cell hyperplasia and worm clearance are both delayed until at least day 14 post infection in *Trpm5^-/-^* mice (Figure 4I-J), although it appears that redundant or compensatory mechanisms are eventually activated. In sum, succinate, *N. brasiliensis*, and *T. muris* all initiate type 2 immune responses through a common sensing mechanism that requires tuft cells and TRPM5.

## Discussion

In this study, we identify succinate as the first intestinal tuft cell ligand. Succinate is sufficient to induce a type 2 immune response in the small intestine, and it does so by activating the tuft-ILC2 circuit through TRPM5 in tuft cells. Previous studies identified tissue-specific tuft cell effector functions, such as smooth muscle contraction in the trachea and urethra, and regulation of the type 2 circuit in the small intestine (Deckmann et al., 2014; Krasteva et al., 2011; von Moltke et al., 2016; Tizzano et al., 2010). Our study now highlights the diversity of tuft cell sensing by providing transcriptional evidence that the ligands and receptors of chemosensing by tuft cells are tissue-specific. Tracheal and thymic tuft cells express canonical taste receptors and, at least in the case of the trachea, can respond to bitter ligands (Krasteva et al., 2011; Tizzano et al., 2010). Tuft cells in the small intestine, on the other hand, are activated by succinate and perhaps by additional taste receptor-independent ligands. It remains to be determined how tuft cells in the gall bladder and colon, which lack expression of both taste receptors and *Sucnr1*, are activated. Recent work has revealed that at least two subsets of intestinal tuft cells exist (Haber et al., 2017); we expect that targeted single cell sequencing of tuft cells in different tissues would likely reveal an even greater diversity of receptors. It will be interesting to see how expression of taste receptors is distributed among tuft cells in the thymus or trachea. Perhaps a small subset of tuft cells in the small intestine also expresses taste receptors but could not be detected in our global analysis. Overall, our findings suggest the intriguing possibility that while all tuft cells share a conserved morphology and express the intracellular machinery for signal transduction, their upstream sensing specificities have evolved to match their microenvironment.

Given that intestinal tuft cells express *Ffar3*, it is perhaps surprising that none of the SCFA ligands for FFAR3 were sufficient to induce a type 2 immune response. It is possible that when SCFAs are given in the drinking water they are rapidly absorbed in the stomach and intestine before they can accumulate to levels that would activate FFAR3. Furthermore, in vitro studies suggest FFAR3 couples principally to Gi/o alpha subunits (Le Poul et al., 2003), in which case signaling through FFAR3 might induce cAMP-regulated non-immune (or even immunoregulatory) tuft cell effector functions that were not detected by our assays here. Whether and how FFAR3 regulates tuft cell biology therefore remains to be determined.

Our discovery of succinate as a type 2 innate ligand also suggests a novel paradigm in which the intestinal type 2 immune response is tuned to microbial metabolism. Helminths, protists, and both commensal and pathogenic bacteria have evolved diverse metabolic strategies to thrive in the nutrient-rich but oxygen-poor intestinal lumen. In many cases, these anaerobic metabolic pathways employ fumarate as the terminal electron acceptor, leading to the production and secretion of succinate. Our findings suggest that this succinate could be sensed by tuft cells to initiate a type 2 immune response. The low frequency of tuft cells in *T. muris-* and specific pathogen-free mice suggests that bacterial succinate is not sensed at homeostasis, but this may change in states of dysbiosis, especially when bacterial species that secrete succinate are expanded.

In this study we have provided initial evidence of a link between *N. brasiliensis* infection and succinate sensing in tuft cells. We have shown that the *N. brasiliensis* exudate contains succinate and can activate SUCNR1, and that the sensing of both succinate and *N. brasiliensis* requires TRPM5, whose expression in the intestinal epithelium is restricted to tuft cells. Thus, it is possible that tuft cells directly sense succinate secreted by *N. brasiliensis* in the intestine. Alternatively, since it is well appreciated that helminth infection alters the microbial composition in the intestine (Fricke et al., 2015; Su et al., 2018), tuft cells may detect *N. brasiliensis* indirectly via modified levels of commensal-derived succinate. Clearly, much more work will be needed to understand how the complex interplay between intestinal helminths, protists and bacteria may result in increased succinate levels that can be sensed by tuft cells.

Experiments using *Sucnr1^-/-^* mice will provide important insights into the role of this pathway in the immune response, but we note accumulating evidence for redundancy in type 2 immune sensing that may mask phenotypes in mice deficient for just one receptor. For example, tuft cell hyperplasia and worm clearance are delayed but do eventually occur in both *Trpm5^-/-^* and *Il25^-/-^* mice (Fallon et al., 2006 and data not shown). Also, *N. brasiliensis* induces tuft cell hyperplasia along the entire small intestine (von Moltke et al., 2016), while the response to succinate is more restricted to the ileum (Figure 3C), indicating that *N. brasiliensis* may activate the tuft-ILC2 circuit through multiple mechanisms.

It should also be noted that we cannot rule out the possibility that tuft cells detect endogenous succinate. In naïve rodents, plasma succinate concentrations range from 5–15 μM and are therefore below the EC50 of 28 μM that has been calculated for murine SUCNR1 (He et al., 2004; Sadagopan et al., 2007), but succinate levels may increase either systemically or locally during infection or stress. Tuft cells may also detect dietary succinate, however, our observation that even 75 mM succinate in the drinking water was insufficient to induce maximal tuft cell hyperplasia suggests that it is difficult to globally raise the lumenal concentration of succinate in the intestine, likely due to rapid transport of succinate across the epithelial barrier (De Vadder et al., 2016; Wolffram et al., 1994). We therefore favor the model that worms and other microbes that gain access to the intestinal epithelium are uniquely suited to locally deliver succinate at levels that are high enough to activate SUCNR1 in tuft cells.

Regardless of its physiologic source, it is interesting to consider succinate in the context of other innate immune ligands. As classically defined, an innate immune ligand should be unique to microbes and difficult to mutate without incurring fitness costs. Succinate, it seems, meets neither of these criteria. First, although normally sequestered inside cells, succinate is abundant in host tissue. Furthermore, although succinate itself is highly conserved, the diversity of metabolic pathways found among helminth species alone suggests that it would not be difficult to evolve away from succinate *secretion* and thereby evade detection by tuft cells. Therefore, we must consider the possibility that protists and helminths can tolerate detection via succinate. In the case of commensal organisms, it is tempting to speculate that some species, such as those that can consume mucus, may even target the succinate receptor to enhance goblet cell mucus production, although the lack of tuft cell hyperplasia in most SPF mice suggests this may only occur with certain microbial compositions. Moreover, since many helminths infect their hosts chronically with only modest losses in fitness for the host, it is possible that these organisms manipulate host sensing pathways to establish a relatively tolerogenic immune state, permitting long-term colonization.

On the host side, why does the immune system tolerate the risk of sensing endogenous succinate? Indeed, in this respect extracellular succinate is more like a danger associated molecular pattern (DAMP), such as ATP or HMGB1, although classically these stimulate inflammatory type 1 responses. Interestingly, the only previous link between SUCNR1 and the immune system is a study showing that signaling through SUCNR1 on dendritic cells can enhance type 1 immune responses (Rubic et al., 2008). Sensing of *intracellular* succinate also enhances type 1 inflammatory signaling (Tannahill et al., 2013).

In the intestines, however, sensing of succinate seems to be wired differently. Perhaps SUCNR1 is restricted to the apical membrane of tuft cells to provide spatial segregation that limits detection of host succinate. In addition, inappropriate activation of a type 2 response presents much less risk to the host than a type 1 response, especially in the intestine. Exogenous activation of Toll-like receptors or inflammasomes is rapidly lethal (Heppner and Weiss, 1965; von Moltke et al., 2012), but the immune response to intestinal succinate appears to be self-limiting and causes little morbidity, in keeping with the nature of type 2 immune responses to microbes. Virtually all wild animals and many humans are chronically parasitized with helminths, and while high worm burdens are undoubtedly detrimental to host fitness, especially in children, lower burdens are well tolerated. Indeed, there is growing evidence that helminth infection can provide therapeutic relief of inflammatory bowel disease and allergies (Helmby, 2015), and a type 2 immune response in adipose tissue protects against metabolic disease (Odegaard et al., 2007; Wu et al., 2011). Our findings therefore also have important therapeutic implications. While systemic manipulation of SUCNR1 signaling is not advisable given its role in physiologic processes such as blood pressure regulation (Ariza et al., 2012), tissue specific targeting of SUCNR1 might be possible with agents that are restricted to the intestinal lumen. In sum, we have uncovered a novel paradigm of metabolite sensing by the type 2 immune response, and therapies that manipulate the underlying tuft-ILC2 circuit may provide access to an immune rheostat that can either enhance anti-helminth immunity or dampen inflammatory disease.

## ACKNOWLEDGEMENTS

We thank J. Jaffe for technical assistance and laboratory support, staff at the University of Washington Gnotobiotic Core for assistance with germ free mouse experiments, C. Schneider, C. Miller and Z. Wang for experimental assistance, N. Arpaia for guidance on NMR analysis of intestinal samples, D. Hailey and the Garvey Cell Imaging Lab in the Institute for Stem & Cell Regenerative Medicine for microscopy support, K. Smith for reagents and advice, and M. Fontana for critical reading and editing of the manuscript. JWM is supported by the University of Washington Immunology Training Grant (T32 AI106677). JVM is a Damon Runyon-Dale Frey Breakthrough Scientist and a Searle Scholar. This work was supported by NIH 1DP2 OD024087 (JVM) and the University of Washington. mRNA sequencing was supported by NIH R01 AI26918 (RML) and the SABRE Center at UC San Francisco.

## AUTHOR CONTRIBUTIONS

MSN and JWM designed and performed experiments, analyzed data, and wrote the paper with JVM. MRLC assisted with experiments. GANG performed NMR analysis, with supervision by DR. JLP performed bioinformatics analysis of mRNA sequencing data, with supervision by DJE. RML acquired funding and provided resources for mRNA sequencing. JVM conceived of and supervised the study, performed experiments, analyzed data, and wrote the paper with MSN and JWM.

---

**Figure 1. RNA-Seq identifies a transcriptional tuft cell signature** Tuft cells (CD45^lo^ EPCAM^+^ Flare25^+^) were sorted from the indicated tissues of naïve B6.*Il25^Flare25/Flare25^* mice for mRNA sequencing. Non-tuft epithelial cells of the small intestine (SI; CD45^lo^ EPCAM^+^ Flare25^-^) were sorted as negative controls. **(A)** Principle component analysis of gene expression data. Each dot represents an individual mouse, except in samples from the trachea where each dot represents cells pooled from four mice. **(B)** Hierarchical clustering of differentially expressed genes among tuft cell subsets with an absolute fold change > 8 (FDR < 0.01). **(C)** Ranked listing of a tuft cell transcriptional signature, comprised of genes that were expressed Log2 fold-change > 4 in all tuft cells relative to non-tuft epithelial cells. **(D)** GO Term enrichment analysis identified a single overrepresented gene category within the tuft cell signature (FDR < 0.05). **(E)** Overrepresented pathways identified by KEGG analysis (FDR < 0.05). See also Figure S1.

**Figure 2. The chemosensory receptor repertoire of tuft cells is tissue-specific (A-C, E, F)** RNA-Seq data generated as described in Figure 1 were analyzed for expression of the indicated genes in tuft cells sorted from small intestine (SI), gall bladder (Gall), colon (Col), trachea (Tra), and thymus (Thy). Non-tuft small intestinal epithelium served as a negative control. **(D)** Heat map depicting expression of all taste receptors in tuft cells from the indicated tissues. **(G)** Tuft cells (Il25^+^) and non-tuft cells (Il25^-^) were sorted from the small intestine of naïve B6.*Il25^Flare25/Flare25^* mice and expression of the indicated genes was assessed by RT-qPCR. A-F show biological replicates from one RNA sequencing experiment. Data in G show biological replicates pooled from three experiments. In A-C and E-G, mean + SEM is shown. In A-C and E-F *, FDR < .05; ***, FDR < .001 by statistical analysis in RNA sequencing pipeline. In G *, p < 0.05; ***, p < 0.001 by multiple t-tests (G). n.s., not significant.

**Figure 3. Succinate is sufficient to induce a type 2 immune response in the small intestine (A-C)** Mice were given the indicated molecules (150 mM, except 300 mM for NaCl) in drinking water for 7 d prior to harvest, sectioning, and labeling of small intestine with anti-DCLK1 (tuft cell label; red) and DAPI (blue). **(A)** Representative images of distal (last 10 cm) small intestine are shown. Scale bar = 50 μm **(B)** Tuft cells were quantified by microscopy. **(C)** Tuft cells were enumerated in the proximal (first 10 cm) and distal (last 10 cm) sections of the small intestine (SI) of mice treated 7 d with 150 mM succinate or a water control. **(D)** Tuft cells were enumerated in the distal (last 10 cm) small intestine following 7 d treatment with the indicated concentrations of succinate. **(E)** Kinetics of tuft cell expansion in distal (last 10 cm) small intestine of mice treated with 150mM succinate (n = 3-9 biological replicates for each timepoint). **(F)** Representative images of middle (10-20 cm from cecum) small intestine stained with periodic acid-Schiff to visualize goblet cell number and hypertrophy following 7 d of 150 mM succinate treatment. Scale bar = 50 μm. Goblet cell numbers **(G)** and hypertrophy **(H)** were quantified by microscopy. **(I)** ILC2 activation was assessed in the lamina propria after 36 hours of 150mM succinate treatment of wild-type or *Trpm5^-/-^* IL-13 reporter mice (Smart13). Representative flow plots are shown. **(J)** Quantification of ILC2 activation. **(K-N)** Mesenteric lymph node cells were collected after 7 d treatment with 150 mM succinate and assessed for ILC2 production of IL-13 **(K, L)** and absolute numbers of ILC2s **(M)** and eosinophils **(N)**. **(O)** Tuft cell expansion was measured in distal (last 10 cm) small intestine of germ-free mice after 7 d treatment with 150 mM succinate. All bar graphs depict mean + SEM of at least two independent experiments except that K-N depict results from one experiment. Each symbol represents an individual mouse. *, p < 0.05; **, p < 0.01; ***, p < 0.001 by one-way ANOVA (B-D, J, L-N) with comparison to control, or by Mann-Whitney (G-H, O). See also Figures S2-3.

**Figure 4. Succinate signals via the tuft cell-ILC2 circuit in a TRPM5-dependent manner (A-C)** Distal (last 10 cm) small intestine was harvested from mice of the indicated genotypes after 7 d treatment with 150 mM succinate and labeled with anti-DCLK1 (tuft cell label; red) and DAPI (blue). **(A)** Representative images are shown. Scale bar = 50 μm **(B)** Tuft cells were quantified by microscopy. **(C)** Representative Alcian blue staining in middle (10-20 cm from cecum) small intestine from tuft cell-replete *(Pou2f3+^/-^*) or tuft cell-deficient (*Pou2f3^-/-^*) mice following 7 d treatment with 150 mM succinate. Scale bar = 100 μm. Goblet cell numbers **(D)** and hypertrophy **(E)** were quantified by microscopy. **(F)** Concentration of the indicated molecules was measured in *N. brasiliensis* excretory-secretory product (NES) or culture media control by NMR. Results shown are representative of three independent experiments. **(G)** Wild-type (B6) or SUCNR1-transduced MEFs were loaded with a fluorescent calcium indicator and fluorescence/background (R/R_0_) was quantified over time. Each gray trace represents one cell. Black line indicates average R/R_0_. Arrows indicate time of addition of test agonist or ionomycin positive control (1ug/mL). _(H)_ The maximal R/R0 of the test agonist was normalized to the R/R0 maximal response to ionomycin for each fluorescent calcium flux sample and quantified. _(I)_ Wild-type and *Trpm5^-/-^* mice were infected with *N. brasiliensis* and tuft cell frequency was assessed in the distal (last 10 cm) small intestine over time. **(J)** Total worm burden in wild-type or TRPM5-deficient mice over the course of *N. brasiliensis* infection. In B, D, and E, each symbol represents one mouse pooled from at least two independent experiments. I and J show the average of 3-10 biological replicates pooled from three independent experiments. G shows representative graphs and H shows technical replicates from more than three independent experiments. All graphs depict mean + SEM. *, p < 0.05; **, p < 0.01; ***, p < 0.001 by one-way ANOVA (B) with comparison to Wt(B6) control, by Mann-Whitney (D-E), or using multiple t-tests (H-J). n.s., not significant; n.d., not detectable.

**Figure S1. Pairwise comparison of differential gene expression** Volcano plots showing pairwise comparisons of differentially expressed genes in the cell subsets analyzed by mRNA sequencing. Red dots represent genes with FDR < 0.01. T_ = tuft cells; E_ = non-tuft epithelium; SI = small intestine; Col = colon; Gall = gall bladder; Thy = thymus; Tr = trachea.

**Figure S2. Gating strategies** Flow cytometry gating strategy for cells from the **(A)** small intestine lamina propria and **(B)** mesenteric lymph nodes.

**Figure S3. Immune cell accumulation in mesenteric lymph node after succinate treatment** Smart13 or *Trpm5-/-;Smart13* mice were either left untreated or given 150 mM succinate in the drinking water for 7 days before mesenteric lymph nodes were harvested and analyzed by flow cytometry. **(A)** Percentage of eosinophils among all live cells. **(B)** Percentage of ILC2s among all live cells. Graphs depict mean + SEM. Each symbol represents an individual mouse and data are from one experiment. **, p < .01; ***, p < .001 by one-way ANOVA with comparison to untreated Smart13 mice.

**Table S1.** Flow cytometry antibodies used in this study

**Table S2.** qPCR primers used in this study

**Item S1.** Differentially expressed genes among all tuft cell subsets

**Item S2.** The transcriptional tuft cell signature

**Item S3.** GO term enrichment analysis of the tuft cell signature

**Item S4.** KEGG pathway enrichment analysis of the tuft cell signature

## MATERIALS & METHODS

### Mice and rats

Mice aged 6 weeks and older were used for all experiments. Mice were age-matched within each experiment, but pooled results include both male and female mice of varying ages. C57BL/6J mice were bred in house or purchased from Jackson Laboratories. *Rag1^-/-^* (B6.129S7-Rag1^tm1M^om/J; 002216), Il2rg^-/-^ (B6.129S4-Il12rg^tm1W^j^hl^/J; 003174), and *Trpm5^-/-^* (B6.129P2-Trpm5^tm1Dgen^/J; 005848) mice were purchased from Jackson Laboratories. *B6.Il25^Flare25/Flare25^* and *B6.Il13^Smart13/Smart13^* mice were generated as previously described (Liang et al., 2012; von Moltke et al., 2016). *Il25^-/-^* mice were generated as previously described (Fallon et al., 2006), generously provided by A. McKenzie, and backcrossed at least 8 generations to C57BL/6J. IL4ra^-/-^ mice were generated as previously described (Mohrs et al., 1999), generously provided by F. Brombacher, and backcrossed at least 8 generations to C57BL/6J. *B6.Pou2f3^-/-^* (Pou2f3 tm1.1(KOMP)Vlcg, Project ID #VG18280) were generously provided by M. Anderson. All mice not purchased from Jackson Laboratories were rederived into the University of Washington vivarium. Lewis rats were purchased from Envigo (LEW/SsNHsd). Except mice noted below, all mice and rats were maintained in specific pathogen-free conditions at the University of Washington and were confirmed to be *Tritrichamonas muris-free* by qPCR. Germ free mice were housed and treated in the University of Washington Gnotobiotic Animal Core. All experimental procedures were approved by the Institutional Animal Care and Use Committee at the University of Washington. B6.*Il25^Flare25/Flare25^* mice used to sort cells for mRNA sequencing were maintained in specific pathogen-free conditions at the University of California, San Francisco. This strain was retrospectively found to be colonized with *Tritrichamonas muris*.

### Short Chain Fatty Acid and Succinate Administration

Mice were provided with sodium succinate hexahydrate (Alfa Aesar), sodium propionate (Arcos Organics), sodium acetate trihydrate (Fisher Chemical), or sodium butyrate (Alfa Aesar) ad libitum in drinking water. Unless otherwise noted, mice were treated for 7 days with 150 mM agonist, except NaCl, which was given at 300 mM to match sodium molarity with succinate treatment.

### Germ Free

C57BL/6J mice were raised under standard germ free conditions in the University of Washington Gnotobiotic Animal Core. For succinate administration, mice were transferred to hermetically-sealed positive pressure isocages as described (Paik et al., 2015) and given autoclaved 150 mM sodium succinate hexahydrate in drinking water ad libitum for 7 days.

### Mouse Infection

Infectious third-stage *N. brasiliensis* larvae (L3) were raised and maintained as described (Liang et al., 2012). Mice were infected subcutaneously at the base of the tail with 500 *N. brasiliensis* L3 and were euthanized at the indicated time points to collect tissues for staining or to count worm burden. Worm burden was enumerated across the entire small intestine. Tuft cells and goblet cells were quantified as described.

### Tissue fixation and staining

For tuft cell staining, intestinal tissues were flushed with PBS and fixed in 4% paraformaldehyde for 4 h at 4° C. Tissues were washed with PBS and incubated in 30% (w/v) sucrose overnight at 4° C. Small intestine and colon samples were then coiled into “Swiss rolls” and embedded in Optimal Cutting Temperature Compound (Tissue-Tek) and sectioned at 8 μm on a Microm HM550 cryostat (Thermo Scientific). Immunofluorescent staining was performed in Tris/NaCl blocking buffer (0.1 M Tris-HCL, 0.15 M NaCl, 5μg ml^-1^ blocking reagents (Perkin Elmer), pH 7.5) as follows: 1 h 5% goat serum, 1 h primary antibody (aDCLK1, Abcam ab31704), 40 min goat anti-rabbit IgG F(ab’)2-AF594 secondary antibody (Life Technologies) and mounted with Vectashield plus DAPI (Vector Laboratories). Tuft cell frequency was calculated using ImageJ software to manually quantify DCLK1+ cells per millimeter of crypt-villus axis. Four 10x images were analyzed for each replicate and at least 30 total villi were counted.

For goblet cell staining, tissues were flushed with PBS, fixed in 10% buffered formalin at 4° C for 3 h, coiled into “Swiss rolls” and returned to formalin. After 24 h tissues were moved to 70% ethanol for storage. Tissue processing, paraffin embedding, sectioning, and staining were performed by the Pathology Research Services Laboratory at the University of Washington. Periodic acid Schiff (PAS) or Alcian blue staining was used to identify goblet cells. Goblet cell frequency was calculated as described above for tuft cells. Hypertrophy was quantified using ImageJ software to measure the area of at least 80 goblet cells for each biological replicate. Brightfield and fluorescent images were acquired on an Axio Observer.A1 inverted microscope (Zeiss).

### Single-cell tissue preparation

For single cell epithelial preparations from small intestines, gall bladder, and colon, tissues were flushed with PBS, opened, and rinsed with PBS to remove intestinal contents. Intestinal tissue was cut into 2-5 cm pieces and incubated rocking at 37° C in 15 mL HBSS (Ca^+2^/Mg^+2^-free) supplemented with 5mM dithiothreitol (DTT, Sigma-Aldrich), 5% fetal calf serum (FCS, VWR), and 10mM HEPES (Gibco). Tissues were vortexed vigorously and supernatant was discarded. Tissues were then incubated rocking at 37° C in 15 mL HBSS (Ca^+2^/Mg^+2^-free) supplemented with 5mM EDTA (Invitrogen), 5% FCS, and 10mM HEPES. Tissues were vortexed thoroughly and released epithelial cells were passed through a 70 μm filter. Tissues were then incubated in fresh EDTA/HBSS solution for 15 minutes, vortexed, and filtered. Supernatants were pooled and washed once before staining for flow cytometry.

For lamina propria preparations small intestinal tissue was processed as above to remove the epithelial fraction. Tissues were then rinsed in 20 mL HBSS (with Ca^+2^/Mg^+2^) supplemented with 5% FCS and 10mM HEPES, shaking at 37° C for 20 minutes. Supernatants were discarded and tissues were incubated in 5 mL HBSS (with Ca^+2^/Mg^+2^) supplemented with 3% FCS, 10mM HEPES, 30 μg mL^-1^ DNase I (Sigma Aldrich), and 0.1 Wunsch mL^-1^ Liberase TM (Sigma Aldrich), shaking at 37° C for 30 minutes. Tissues were vortexed and cells were passed through a 70 μm filter and washed. The resulting pellet was resuspended in 40% Percoll (Sigma Aldrich), centrifuged for 5 minutes at 1500 rpm, and supernatant discarded. Pelleted cells were then washed and stained for flow cytometry.

For mesenteric lymph node preparations, MLN were harvested into RPMI + 5% FBS on ice. Tissues were incubated 30 minutes at 37° C in 5 mL RPMI supplemented with 2 mg mL^-1^ collagenase VIII (Sigma-Aldrich) and 7.5 μg mL^-1^ DNaseI (Sigma-Aldrich). Digested MLN were passed through a 70 μM filter and remaining tissue was mashed through filter. Cells were washed once and stained for flow cytometry.

For tracheal epithelium, the trachea was dissected from mice, stored in DMEM + 5% FBS and cleaned of stroma using a dissecting microscope. Tissue was cut into 6 pieces and incubated at room temperature in 16U mL^-1^ Dispase (Corning) in DPBS without Ca^2+^/Mg^2+^. Incubation was monitored carefully and stopped by transfer into DMEM + 5% FBS when epithelium began to lift away from stroma (23-28 minutes). Epithelium was then peeled off under a dissecting microscope and collected in DMEM + 5% FBS on ice. Harvested epithelium was digested in 0.5% Trypsin + EDTA for 20 minutes at 37° C. After digest, cells were vortexed briefly and trypsin was quenched with DMEM + 5% FBS. Cells were washed once and stained for flow cytometry.

For thymic epithelium, thymi were isolated, cleaned of fat and transferred to DMEM + 2% FBS on ice. Thymi were minced with a razor blade and tissue pieces were moved with a glass Pasteur pipette to 15 ml tubes and vortexed briefly in 10 ml of media. Fragments were allowed to settle before removing the media and replacing it with 4 ml of digestion media containing 2% FBS, 100 μg mL^-1^ DNase I (Roche), and 100 μg mL^-1^ Liberase TM (Sigma-Aldrich) in DMEM. Tubes were moved to a 37° C water bath and fragments were triturated through a glass Pasteur pipette at 0 min and 6 min to mechanically aid digestion. At 12 min, tubes were spun briefly to pellet undigested fragments and the supernatant was moved to 20 ml of 0.5% BSA (Sigma-Aldrich), 2 mM EDTA (TekNova), in PBS (MACS buffer) on ice to stop the enzymatic digestion. This was repeated twice for a total of three 12 min digestion cycles, or until there were no remaining tissue fragments. The single cell suspension was then pelleted and washed once in MACS Buffer. Density-gradient centrifugation using a three-layer Percoll gradient (GE Healthcare) with specific gravities of 1.115, 1.065, and 1.0 was used to enrich for stromal cells. Cells isolated from the Percoll-light fraction, between the 1.065 and 1.0 layers, were then washed and resuspended for staining.

### Flow cytometry and cell sorting

Single cell suspensions were prepared as described and stained in DPBS + 3% FBS with antibodies to surface markers for 20 min at 4° C, followed by DAPI (Roche) for dead cell exclusion. See Table S1 for a list of antibodies used in this study. Samples were FSC-A/SSC-A gated to exclude debris, SSC-H/SSC-W gated to select single cells and gated to exclude DAPI+ dead cells. Samples were run on an LSR II (BD Biosciences) or Aurora (Cytek) and analyzed with FlowJo 10 (Tree Star). For cell sorting, single cell suspensions were prepared and stained as described and sorted into CD45^kl^ IL25(RFP)^+^ EpCAM^+^ and CD45^kl^ IL25(RFP^-^) EpCAM^+^ populations using a MoFlo XDP (Beckman Coulter) or an Aria II (BD Biosciences).

### RNA sequencing & analysis

Single cell suspensions of epithelial cells from gall bladder, small intestine, colon, thymus, and trachea were generated as described above from *Il25^Flare25/Flare25^* reporter mice. CD45^lo^ RFP^+^ EpCAM^+^ tuft cells were sorted from all tissues and CD45^lo^RFP^-^EpCAM^+^ cells were also sorted from the small intestine. With the exception of tracheal tuft cells, which were pooled from four mice for each replicate, each biological replicate represents one mouse. Four biological replicates were collected for each sample. Average sorted cells for each sample were as follows: SI_RFP+: 35,250; SI_RFP^-^: 55,000; Colon: 14,775; Gall: 2287, Thymus: 1612; Trachea: 255.

Cells were sorted directly into lysis buffer from the Dynabead mRNA Direct Purification Kit (ThermoFisher) and mRNA was isolated according to the manufacturer’s protocol. Amplified cDNA was prepared using the NuGen Ovation RNA-Seq system V2 kit, according to the manufacturer’s protocol (NuGen Technologies). Sequencing libraries were generated using the Nextera XT library preparation kit with multiplexing primers, according to manufacturer’s protocol (Illumina). Library fragment size distributions were assessed using the Bioanalyzer 2100 and the DNA high-sensitivity chip (Agilent Technologies). Library sequence quality was assessed by sequencing single-end 50 base pair reads using the Illumina MiSeq platform and were pooled for high-throughput sequencing on the Illumina HiSeq 4000 by using equal numbers of uniquely mapped reads (Illumina). Twelve samples per lane were multiplexed to ensure adequate depth of coverage. Sequencing yielded a median read depth of 89.2 million reads per sample. The analytic pipeline included de-multiplexing raw sequencing results, trimming adapter sequences, and aligning to the reference genome. Sequence alignment and splice junction estimation was performed using the STAR (https://code.google.com/p/rna-star) software program. For differential expression testing, the genomic alignments were restricted to those that map uniquely to the set of known Ensembl IDs (including all protein coding mRNAs and other coding and noncoding RNAs). STAR aggregated mappings on a per-gene basis were used as raw input for normalization by DESeq2 software. Replicates failing quality control at any stage were discarded.

To determine the tuft cell signature, any genes with a mean normalized read count <300 in any tuft cell subset were removed from analysis. Next, genes were ranked based on log2 fold change between the mean expression level in all tuft samples and the mean expression level in all non-tuft samples, with a cutoff of 4. GO Term and KEGG pathway analysis was performed using the Database for Annotation, Visualization, and Integrated Discovery (Huang et al., 2009a, 2009b). Visualization of taste receptor expression was generated using Morpheus (https://software.broadinstitute.org/morpheus/).

### Quantitative RT-PCR

CD45^lo^RFP^+^EpCAM^+^ and CD45^lo^RFP^-^EPCAM+ populations from *Il25^Flare25/Flare25^* mice were sorted into Buffer RLT (Qiagen) as described. RNA was isolated using the Micro Plus RNeasy kit (Qiagen) and reverse transcribed using SuperScript Vilo Master Mix (Life Technologies). The resulting cDNA was used as a template for quantitative PCR with Power SYBR Green reagent on a StepOnePlus cycler (Applied Biosystems). Transcripts were normalized to *Rps17* (40S ribosomal protein S17) expression. See Table S2 for a list of primers used in this study.

### Nuclear Magnetic Resonance Quantification of SCFA and Succinate

NMR analyses were made using a Bruker AVANCE III 800 MHz equipped with a 5 mm HCN cryoprobe suitable for ^1^H inverse detection with Z-gradients at 298 K. Samples were prepared to contain 50 μM TSP (3-(trimethylsilyl) propionic-2,2,3,3-d4 acid sodium salt) for quantitative and chemical shift reference. One-dimensional ^1^H NMR spectra were obtained using the CPMG (Carr-Purcell-Meiboom-Gill) pulse sequence that included residual water signal suppression from a pre-saturation pulse during the relaxation delay. For each spectrum, 32k data points were acquired using a spectral width of 9615 Hz. The raw data were processed using a spectral size of 32k points and by multiplying with an exponential window function equivalent to a line broadening of 0.3 Hz. The resulting spectra were phase and baseline corrected and referenced with respect to the internal TSP signal. Bruker Topspin version 3.5pl6 software package was used for NMR data acquisition and processing. Assignment of peaks was made based on ^1^H NMR chemical shifts and spin-spin couplings obtained from the spectra of standard compounds under similar experimental conditions at 800 MHz. Chenomx NMR Suite Professional Software package (version 5.1; Chenomx Inc., Edmonton, Alberta, Canada) was used to quantitate metabolites. This software allows fitting spectral lines using the standard metabolite library for 800 MHz ^1^H NMR spectra and the determination of concentrations. Peak fitting with reference to the internal TSP signal enabled the determination of absolute concentrations for the short chain fatty acids and other organic acids. All NMR experiments were performed in conjunction with the Northwest Metabolomics Research Center at the University of Washington.

### Preparation of *N. brasiliensis* excretory-secretory product (NES)

Infectious third-stage *N. brasiliensis* larvae (L3) were raised and maintained as described. Lewis rats were infected subcutaneously with 2000 *N. brasiliensis* L3. Mature (L5) worms were collected from the entire small intestine 7 days post infection. Worms were washed 10 times in Wash Solution I (PBS with 200U ml^-1^ Pen-Strep), allowing worms to settle by gravity between each wash. Worms were allowed to equilibrate in Wash Solution II (RPMI 1640 with 200U ml^-1^ Pen-Strep) for 1 hour at room temperature, before being transferred to a tissue culture flask in NES culturing media (RPMI 1640, with 100U ml^-1^ Pen-Strep, 2mM L-Glutamine and 1% glucose) and cultured at 37° C. Supernatant was collected at 24 and 48 hours and filtered prior to use as NES. For NMR analysis, phosphate buffer prepared in deuterated water (0.1M; pH =7.4) containing TSP (3-(trimethylsilyl) propionic-2,2,3,3-d4 acid sodium salt) was added to NES to achieve a final concentration of 50uM TSP.

### Cell culture and transduction

Immortalized C57BL/6J mouse embryonic fibroblasts (MEFs; kindly provided by D. Stetson) and HEK293T (Cat# CRL-3216, ATCC) were cultured in DMEM supplemented with 10% fetal calf serum (VWR), L-glutamine, penicillin/streptomycin, sodium pyruvate, HEPES and β-mercaptoethanol. For lentiviral transduction, the ORF for mouse *Sucnr1* was obtained directly from prepared cDNA from sorted small intestinal tuft cells (described above). After sequencing verification, *mSucnr1* was cloned into the pRRL lentiviral vector downstream of an MND promoter as previously described (Gray et al., 2016). VSV-G pseudotyped, self-inactivating lentivirus was prepared by transfecting HEK293T cells with 1.5 μg pVSV-G, 3 μg psPAX-2, and 6 μg pRRL lentiviral vector, and viral supernatant collected and filtered before use. MEFs were incubated with viral supernatant overnight, and media was replaced with fresh culture media. Puromycin (5ug mL^-1^) selection was added 72 hours post transduction and continued for 7 days.

### Calcium imaging

Mouse embryonic fibroblasts were plated 7 × 10^4^ cells/well in 24-well plates coated with poly-D-lysine. After overnight incubation, cells were washed with assay buffer (1X HBSS with Ca^2+^/Mg^2+^, 10mM HEPES, pH 7.4). Cells were incubated for 1 hr at 37°C in assay buffer supplemented with 2.5 mM Fluo-4AM (Invitrogen) and 0.05% pluronic-F127 (Invitrogen). Cells were washed twice with assay buffer and incubated in assay buffer with 1mM probenecid (Biotium) for 30 minutes at 37° C prior to imaging. Cells were maintained at 37° C with 5% CO2 throughout imaging. While imaging, sodium succinate was added to assay buffer at a final concentration of 150uM or 100ul NES or NES culturing media was added to 250ul assay buffer. Following addition of test agonist, ionomycin was added at a final concentration of 1ug mL^-1^. Fluorescence images were collected at 1.44 frames per second with a 40X extra-long working distance objective on a Nikon TiE Inverted Widefield Fluorescence Nikon microscope and analyzed with NIS Elements AR 3.2 software. More than 50 cells were analyzed per replicate.

### Statistical analysis

All experiments were performed using randomly assigned mice without investigator blinding. All data points and “n” values reflect biological replicates (i.e. mice), except in 4H where they represent technical replicates. No data were excluded. Statistical analysis was performed as noted in figure legends using Prism 7 (GraphPad) software. Holm-Sidak was used to correct for multiple comparisons. Graphs show mean + SEM.

